# Subthreshold Asynchronous States and Computations in Biophysically Detailed Populations of Neurons

**DOI:** 10.64898/2026.04.22.720183

**Authors:** Wilten Nicola, Ali Mahani, Raymond W. Turner, Gerald Zamponi

## Abstract

Spikes are metabolically costly to generate and transmit, and spiking rates should be kept to a minimum for efficient coding. Many invertebrate circuits do not fire spikes or fire spikes at extremely low rates by using graded synaptic transmission as the communication mechanism between neurons. We show that computations and asynchronous states based on excitatory/inhibitory balance can exist without firing any spikes through self-sustaining subthreshold voltage fluctuations powered by graded synaptic transmission in networks of biophysically detailed Hodgkin-Huxley neurons. This novel subthreshold asynchronous state, which we call subthreshold voltage chaos, can be controlled for useful computation and pattern generation, also without firing spikes. Further, we identify candidate ion channels, low-voltage-activated T-Type calcium channels that provide a biophysical mechanism for this type of subthreshold computation. Our work demonstrates a mechanism for computation with graded synaptic release with very few spikes.

## Introduction

The efficient coding hypothesis, originally proposed by Horace Barlow in 1961 [1], postulates that neural circuits should minimize the number of excess spikes fired. From an evolutionary perspective, this hypothesis is reasonable as spikes are metabolically costly to generate and propagate across the synaptic cleft [2].

Indeed, many invertebrate circuits do not fire spikes or fire spikes at extremely low rates. For example, the classical lobster stomatogastric ganglion (STG) utilizes graded release [3, 4] to sustain rhythmic oscillations [5]. Crustaceans also use similar graded-release powered central pattern generators to control swimming [6, 7]. The graded release of neurotransmitter has also been documented in the locust [8], nematode [9], and more broadly, plays a role in retinal neurons [10].

In such a subthreshold regime, neurons would have to perform recurrent computations and generate patterns without firing spikes [11, 12]. This type of fully analog computation may be mediated by low-voltage activated calcium channels like the T-Type channel [13–16] which can support subthreshold computation through the existence of window currents that only operate in voltage regimes near the resting membrane potential [13, 14], and cause synaptic vesicle exocytosis in the absence of spikes [17]. Can these channels power analog computations without spikes as seen in invertebrate circuits?

By using biophysically detailed Hodgkin-and-Huxley neurons coupled with conductance based synapses, we demonstrate rich subthreshold dynamics that mimic the classic asynchronous states that emerge under excitatory/inhibitory balance in rate networks. This novel asynchronous state, which we call subthreshold voltage chaos emerges when neurons have graded synaptic release at voltages near the resting membrane potential. Subthreshold voltage chaos is self-sustaining without firing spikes when the conductances are suitably weak, but can generate sparse spiking for stronger conductances which can also be used to convey information. Further, by using First Order Reduced and Controlled Error (FORCE) training [18–21], we demonstrate that voltage chaos can be tamed for subthreshold pattern generation and computation. Moreover, voltage chaos can be mediated by T-type calcium channels, which lead to calcium influx in the presynaptic axon at voltages near the resting membrane potential. Our results provide computational evidence for spike-independent or silent computation [22], where neural circuits can operate in an exclusively analog regime, with the spikes only used to digitize and transmit the analog computation from one circuit to the next.

## Results

Many neural circuits perform recurrent computations with spikes (Figure 1A). The spikes are transmitted and actively take part in recurrent loops, either locally in brain areas like the hippocampus [23], or in more extended recurrent networks involving multiple brain circuits. These circuits perform computations digitally, either in part or entirely, and transmit information digitally with spikes. Here, we propose an alternative but complimentary mechanism where neural circuits consisting of neurons with tight coupling between somatic, dendritic, and axonal compartments (Figure 1B) instead perform computations in an analog manner through their subthreshold dynamics [11, 12, 24, 25]. Instead of communicating recurrently with full synaptic release triggered from spikes, they smoothly transduce their voltage fluctuations into smaller, so-called “graded” neurotransmitter release [24, 25]. These neurons may be small, or potentially larger and insulated by myelin sheath.

**Figure 1.**
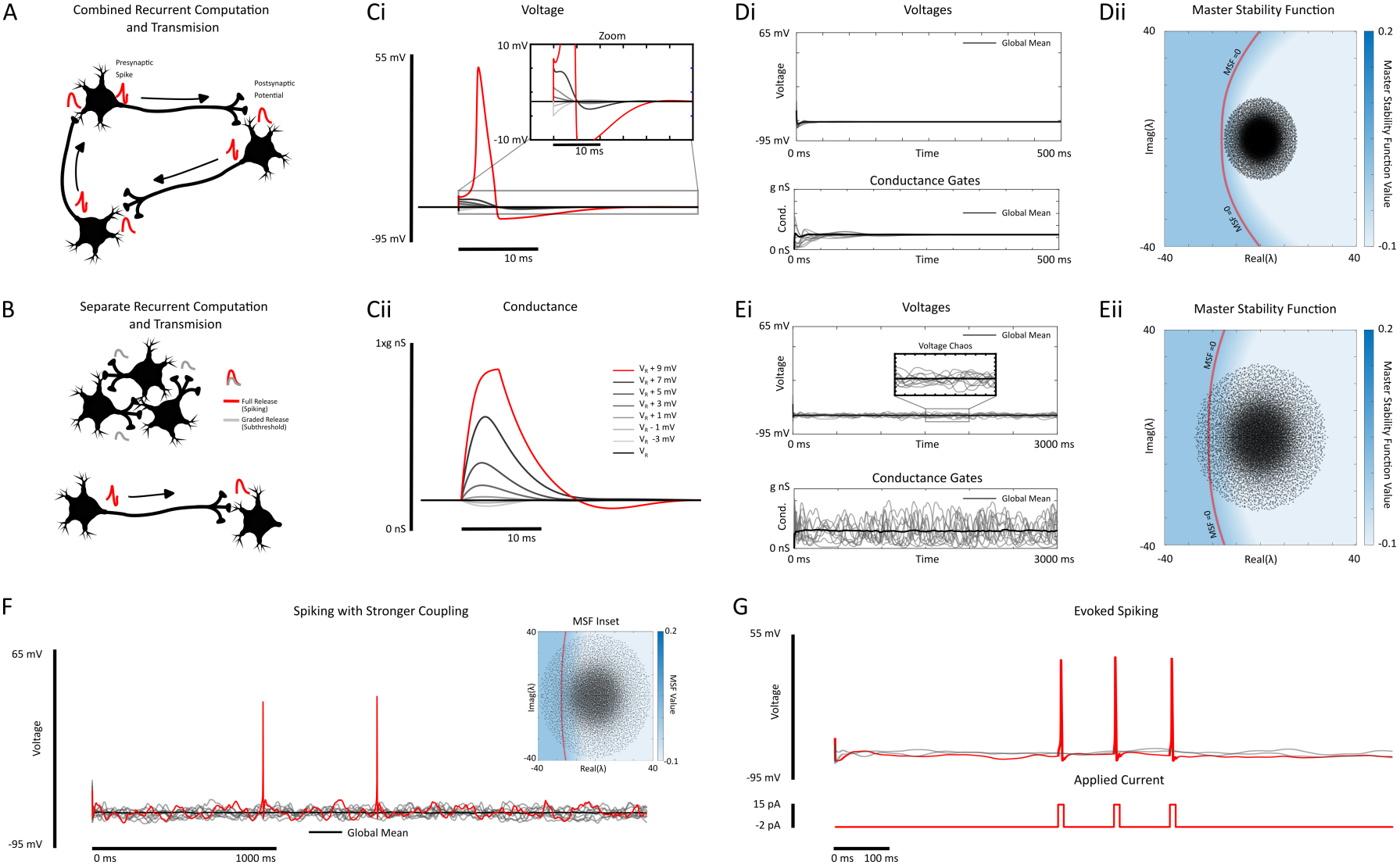
Graded synaptic release powers subthreshold asynchronous states. **(A)** In conventional models of recurrent computation, neurons fire spikes in recurrent loops. The spikes produce postsynaptic potentials and more spikes. **(B)** Here, we consider separate recurrent computation powered by graded release, with spikes used to transmit information, rather than taking part in the recurrent computation directly. **(C)**(i) A single Hodgkin-Huxley neuron is initialized at 7 different membrane potentials from *V*_*R*_ − 3 mV to *V*_*R*_ + 9 mV in increments of 2 mV. A spike is generated only for *V*_*r*_ + 9 mV. (ii) The resulting post-synaptic conductance exhibits graded levels as a function of the presynaptic voltage. **(D)**(i) The voltages (top) and conductances for 10 neurons (grey) and the network means (black) with 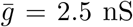. (ii) The master stability function (MSF) values (blue) and the eigenvalues (black dots) for the weight matrix from Di. The red-dashed line corresponds to the MSF change sign, resulting in a loss of stability for the resting membrane potential. **(E)** Identical as in (D) only with 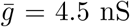. **(F)** An identical network as in (D)-(E), only with 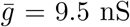 eliciting sparse spikes. A single spiking neuron is highlighted in red. **(G)** An externally elicited spike with 15 pA applied current pulses for a neuron where from the network considered in (E). Parameters for the network under consideration can be found in Table 1.

We investigated the potential for analog computation by considering networks of biophysically detailed Hodgkin and Huxley neuron models [26, 27] with conductance-based coupling [28]:

**Table 1.** The parameters used for the Hodgkin-Huxley and Krinskii-Kokoz neuron model for all figures. The parameter label, the description, and the value + units are shown in the 1st, 2nd, and 3rd column. Note that following Hodgkin and Huxley’s original convention, we write down the parameters assuming that the resting membrane potential has been offset to 0 mV, with all parameters measured relative to that. All figures however are reported without offset to aid in reader interpretability. The membrane capacitance was increased for the Krinskii-Kokoz as we empirically found this was qualitatively closer to the subthreshold voltage-chaos regime in the full Hodgkin-Huxley model. The *V*_1*/*2_ and *K*_*p*_ parameters for Figure 4 (inactivation/activation gates) are reported in the Materials and Methods.

| Parameter | Parameter Description | Value and Units |
|---|---|---|
| *C* | Membrane Capacitance | 1 *µ*F/cm<sup>2</sup> (10 *µ*F/cm<sup>2</sup> for Krinskii-Kokoz) |
| *I* | Background Current | -2 pA |
| *V*<sub>*Na*</sub> | Sodium Reversal Potential | 120 mV |
| *V*<sub>*K*</sub> | Potassium Reversal Potential | -12 mV |
| *V*<sub>*L*</sub> | Leak Reversal Potential | 10.613 mV |
| *g*<sub>*Na*</sub> | Sodium Unitary Conductance | 120 mS/cm<sup>2</sup> |
| *g*<sub>*K*</sub> | Potassium Unitary Conductance | 36 120 mS/cm<sup>2</sup> |
| *g*<sub>*L*</sub> | Leak Conductance | 0.3 120 mS/cm<sup>2</sup> |
| *T*<sub>*max*</sub> | Max Neurotransmitter Release | 1 (unitless) |
| *a*<sub>*r*</sub> | Synaptic Rise Parameter | 1.1 ms<sup>−1</sup> |
| *a*<sub>*d*</sub> | Synaptic Decay Parameter | 0.1 ms<sup>−1</sup> (0.18 ms<sup>−1</sup> Figure 1C) |
| *V*<sub>1/2</sub> | Half-Max Release Voltage | 5 mV |
| *K*<sub>*p*</sub> | Sensitivity Parameter | 2 mV |
| *E*<sub>*gaba*</sub> | GABA Reversal Potential | -20 mV |
| *E*<sub>*ampa*</sub> | AMPA Reversal Potential | 65 mV |

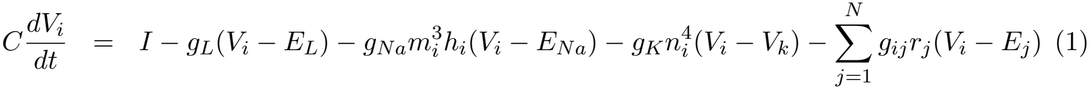

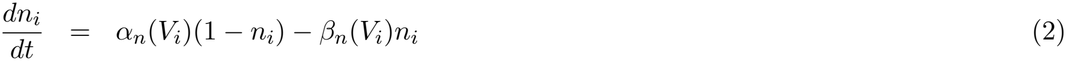

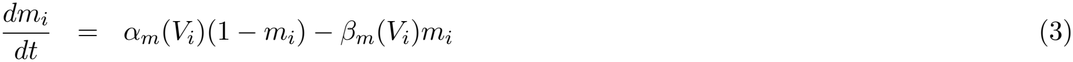

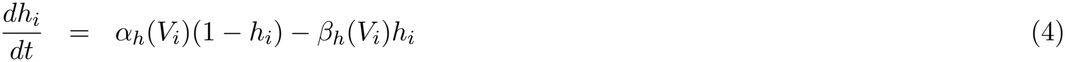

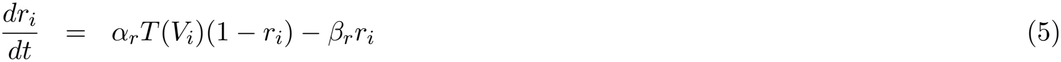

where *V*_*i*_(*t*) corresponds to the voltage of the *i*th neuron, and *m*_*i*_, *h*_*i*_, *n*_*i*_ correspond to the sodium activation gate, sodium inactivation gate, and potassium activation gate, respectively. All neuronal parameters are the standard Hodgkin-Huxley parameters originally fit (Materials and Methods, Table 1) with slight modifications to rectify the voltages and shift the single neuron’s resting membrane potential to −65 mV. The conductance gate *r*_*i*_ coupled neuron *i* neuron *j* via *g*_*ij*_*r*_*j*_(*V*_*i*_ − *E*_*j*_) where *E*_*j*_ acts as the reversal potential for the synapses formed by neuron *j*, and *E*_*j*_ = *E*_*ampa*_ for excitatory neurons, and *E*_*j*_ = *E*_*gaba*_ for inhibitory neurons.

The function *T* (*V*) couples the presynaptic voltage to postsynaptic conductance changes, and is given by

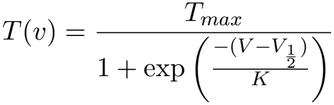

The coupling function *T* (*v*) has been used extensively in computational neuroscience, having been previously derived in [28] on the assumption that the cascade of events leading to changes in the postsynaptic conductance (e.g. presynaptic Calcium release, vesicle fusion, neurotransmitter drift, etc.) are fast enough to be regarded as instantaneous. We have not created a novel type of synaptic connection, but are using existing models of the biophysics behind synaptic transmission.

However, we make one assumption on the parameters for *T* (*V*): we assume that the threshold for neurotransmitter release, given by the half-max release parameter 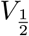 is slightly more depolarized (by 5-10 mVs) than a cell’s resting membrane potential *V*_*R*_. This modification alone is enough to trigger robust graded synaptic release (Figure 1C) as demonstrated by clamping a single neuron to different initial states, and allowing a return to the resting state. Small increases or decreases to the postsynaptic conductance can occur in response to different depolarized or hyperpolarized presynaptic voltages (Figure 1C). As we show later, this change also mimics some of the dynamics of T-type calcium channels.

### The Emergence of Subthreshold Voltage Chaos

With this simple model of graded release, we sought to determine if a network of biophysically detailed neurons could produce rich non-resting subthreshold dynamics powered by graded release. The theory of excitatory/inhibitory balanced networks provides a recipe for how this should be achieved [29–37]. In particular, the excitatory and inhibitory connections should balance with mean 0, and the weights should scale like 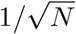 , where *N* is the network size. In order to apply the theory of balanced networks, we require three conditions on the conductances *g*_*ij*_ (Materials and Methods)

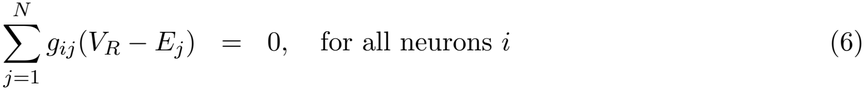

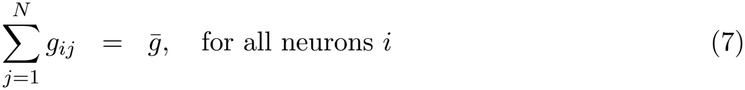

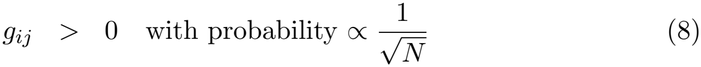

Condition (6) implies all neurons fluctuate around the common resting membrane potential *V*_*i*_ = *V*_*R*_ at steady state. Further, the effective “weight” coupling neuron *j* to neuron *i, ω*_*ij*_ is also given by *ω*_*ij*_ = *g*_*ij*_(*V*_*R*_ − *E*_*J*_), and so condition (6) also implies that the excitatory (*ω*_*ij*_ *<* 0) and inhibitory weights (*ω*_*ij*_ *>* 0) balance each other. Note that the weight signs are reversed from the normal convention as an additional negative sign appears in applying the synaptic currents. Condition (7) is required to analytically determine the stability of the common resting membrane potential *V*_*i*_ = *V*_*R*_ while condition (8) and (7) together lead to the remaining conductances (and effective weights) to scale like 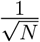 (Materials and Methods). Collectively, the three assumptions on the conductances can be interpreted as the network being sparsely coupled with strong connections that balance excitation and inhibition.

With the conditions on the conductances determined, we sought to determine if rich dynamics would emerge, and if the rich dynamics could be confined to the subthreshold regime. This is critical to subsequently use these rich dynamics as a basis for computation [18–21].

We considered networks where all neurons receive a small hyperpolarizing subthreshold applied current *I* (Figure 1D, Materials and Methods). For a small total conductance 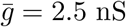, the resting membrane potential was stable (Figure 1Di). This was confirmed by computing the eigenvalues of the steady state resting membrane potential with the master stability function (Figure 1Dii, Materials and Methods, [38–42]). In the master-stability-function framework, the eigenvalues of synchronized solutions are directly determined by the eigenvalues of the weight matrix by the master stability function if the weights are precisely balanced 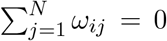, Materials and Methods). We verified that all eigenvalues of the effective weight matrix ***ω*** lead to a stable common resting state for all neurons as the master stability function was strictly negative for all eigenvalues of ***ω*** (Figure 1Dii).

Once 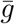 was increased further, the weight matrix eigenvalues crossed the zero-contour of the master stability function (Figure 1E), implying a loss of stability of the stable resting state. This lead to self-sustaining irregular voltage and conductance fluctuations which we call voltage chaos (Figure 1Ei). For 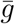 large enough to destabilize *V*_*i*_ = *V*_*R*_, but not too large, the voltage fluctuations did not lead to spikes in any neuron in the network 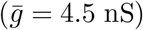. Larger values of 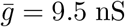 could lead to sparse spikes being fired (Figure 1F, 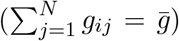). Further, we verified that neurons were still capable of firing spikes when an external current was also applied while the network was in the subthreshold chaotic regime (Figure 1g), indicating that spikes weren’t blocked from firing. Rather, voltage fluctuations around the resting membrane potential *V*_*R*_ but below the threshold to fire spikes are generated through graded depolarization.

Finally, we remark that the constant row-sum assumption 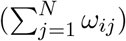 and the precise balance assumption 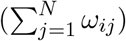,while mathematically convenient for discovering the parameter regimes where voltage fluctuations emerge, they are not necessary for generating these voltage fluctuations (Supplementary Figure 1). In particular, conductances matrices were generated without these conditions and voltage fluctuations (without spiking) still readily emerged, with each neuron having a different resting membrane potential that destabilizes for larger conductances (Supplementary Figure 1).

### A Simpler Interpretable Model of Voltage Chaos

With the existence of rich subthreshold dynamics confirmed in Hodgkin-Huxley networks, we sought to determine if a simple interpretable model could be constructed that would confirm that the subthreshold voltage chaos in Figure 1 is similar to the classical asynchronous states observed in [29–31, 34, 35] shown in both recurrent neural networks [29–31] and in spiking/bursting integrate-and-fire networks [34, 35], with the key difference being the spike-less nature of sub-threshold voltage chaos. In particular, we sought to link the subthreshold dynamics to existing models that display asynchronous states [30, 31, 33].

First, we considered a simpler 2D neuron model: the Krinskii-Kokoz reduction to the Hodgkin-Huxley neuron [27]:

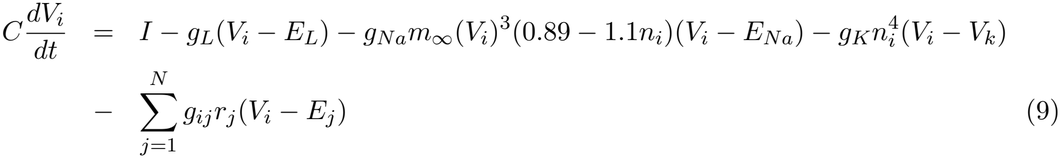

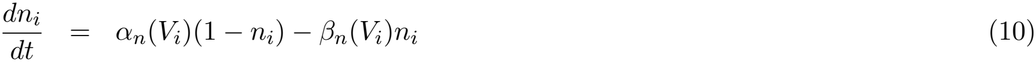

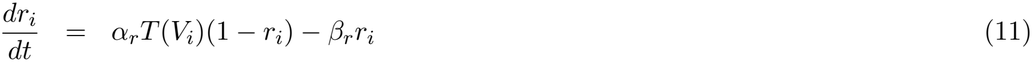

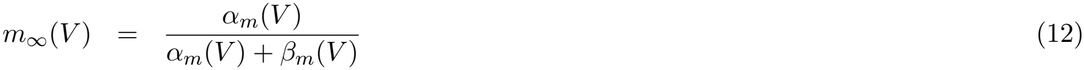

This reduces the system down to a 3*N* dimensional system of differential equations. Next, we assume that the synaptic time constants are fast (AMPA/GABA_A_, ≈ 10 ms) and that the gating variable *r*_*i*_(*t*) is well approximated by its steady state:

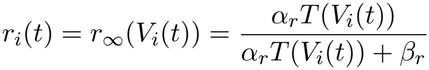

which resulted in a simpler system consisting of 2*N* equations describing a coupled network of conductance neurons with fast-conductance based synapses:

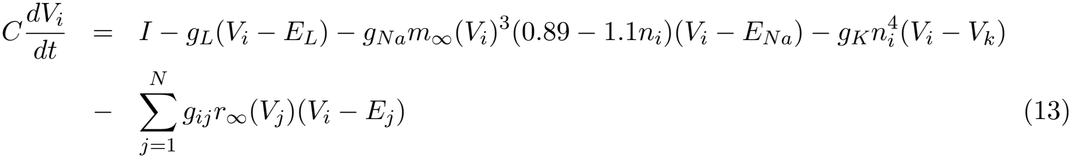

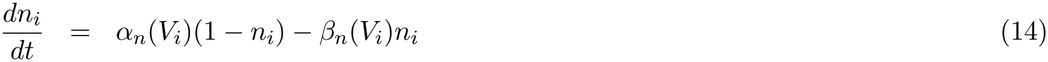

Next, we confirmed the existence of voltage chaos for the Krinskii-Kokoz network by analytically computing the Master Stability Function for the Krinskii-Kokoz network’s global resting membrane state (*V*_*i*_ = *V*_*R*_, Materials and Methods), and simulating 3 networks with a stable resting membrane state (Figure 2Ai), subthreshold voltage chaos at the onset of instability (Figure 2Aii), and sparse spiking for larger conductances (Figure 2Aiii).

**Figure 2.**
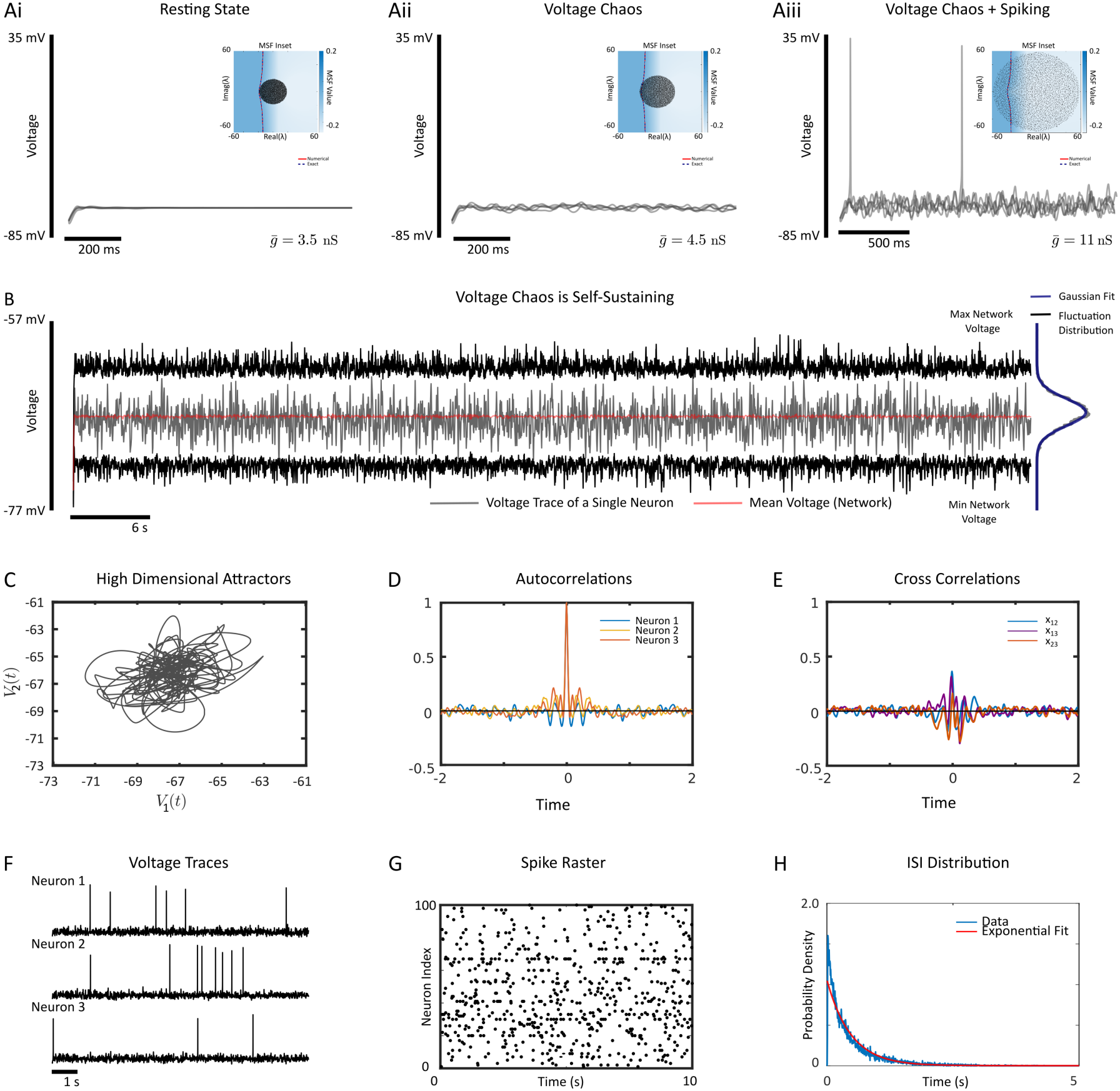
A 2D conductance model shows that subthreshold voltage chaos is an asynchronous state. **(A)** The voltage traces for 5 neurons are shown for 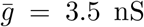 (left), 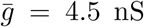 (middle), and 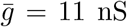 (right). (Inset) The master stability function (blue, numerical, red-dashed, analytic) and eigenvalue spectrum for the network of Krinskii-Kokoz neurons. **(B)** A long simulation of a Krinskii-Kokoz network coupled with 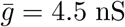 with a single voltage trace shown in grey. The network average (red), maximum network voltage (black, top) and minimum network voltage (black, bottom) are shown. No spikes were observed for 50 seconds of simulation time. The distribution of voltage fluctuations (black, right) are well modeled by a Gaussian distribution (blue, right). **(C)** A plot of *V*_1_(*t*) vs *V*_2_(*t*) showing irregular fluctuations and a high-dimensional attractor for 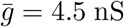. **(D)** The autocorrelation function for 3 neurons in the network. **(E)** The cross correlation for 3 neuron pairs in the network. **(F)** The voltage traces for 3 neurons with 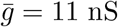 nS, which is sufficient to elicit spiking. **(G)** The spike raster-plot for 100 neurons in the network for 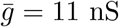. **(H)** The inter-spike-interval (ISI) distribution measured from a 10 second simulation (red) with an exponential fit (red) for 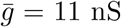. Parameters can be found in Table 1.

With the existence of voltage chaos confirmed in the Krinskii-Kokoz network, we considered the fluctuations in more detail by expanding out the voltages and conductance n-gates as:

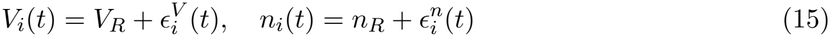

where 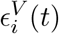 and 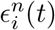 are the time-varying fluctuations around the common resting state *V*_*i*_ = *V*_*R*_ and *n*_*i*_ = *n*_*R*_ that all neurons had. This yielded the following closed-form partially linearized

system which approximated the voltage and *n*-gate fluctuations:

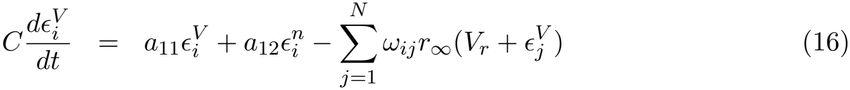

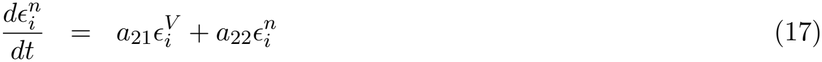

where the coefficients *a*_11_, *a*_12_, *a*_21_, *a*_22_ are derived in the Materials and Methods and *ω*_*ij*_ = *g*_*ij*_(*V*_*R*_ − *Ej*). Note that we have not linearized *r*_*∞*_(*V*_*i*_), as the fluctuations coupled with the low-voltage threshold make this term highly nonlinear. Through a rescaling of time, and the reparameterization of equations (114)-(115), the system of fluctuations is transformed to:

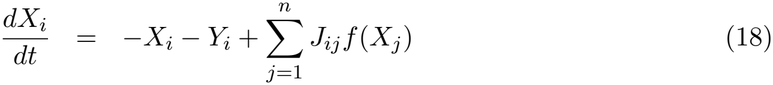

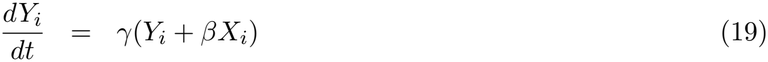

which, for suitable choices of the synaptic weights, *J*_*ij*_, is precisely the recurrent neural network studied in [33]. Notably, this network exhibits a classical asynchronous state, only with resonant dynamics which can be observed in neuronal cross and auto-correlation functions (see Figure 2a in [33]). This derivation shows that the subthreshold dynamics mimic a type of high-dimensional chaotic asynchronous state, where the neuronal parameters would dictate if the chaos is resonant or more broadband.

To test the theory, we simulated the Krinskii-Kokoz network with fast synapses for 50 seconds (Figure 2B). For the entire simulation, no neuron fired a single spike, as shown by the maximum of network voltage fluctuations (Figure 2B), indicating that the subthreshold voltage chaos is self-sustaining. Next, we found that as predicted in [33] and more broadly classical theories of asynchronous states, the fluctuations are well described by a Gaussian distribution (Figure 2B), with high-dimensional attractors (Figure 2C), and voltage auto-correlations displaying damped oscillations (Figure 2D), and weak cross-correlations (Figure 2E). Further, for stronger conductances, neurons sparsely fired spikes (Figure 2F-G) with irregular inter-spike-intervals drawn from a distribution well approximated by an exponential fit (Figure 2H), indicating Poisson-like firing [34, 35, 43]

Collectively, these results support the idea that the voltage fluctuations mimic the classical asynchronous states first suggested in [29–31] only with the potential for resonant dynamics in the subthreshold regime in more recent work [33]. This establishes the existence of subthreshold recurrent neural network (RNN) dynamics powered by graded depolarization. Unlike classical models that link RNNs to spiking rates, the RNNs here operate through subthreshold voltage dynamics without spikes.

### Taming Voltage Chaos for Recurrent Computation with FORCE

With rich subthreshold dynamics confirmed, we next sought to investigate if voltage chaos could be “tamed” for subthreshold computations. As the subthreshold dynamics mimic a recurrent neural network or a reservoir, there are many options in training the effective weights [18– 21, 44, 45]. We utilized the First Order Reduced and Controlled Error (or FORCE, [18, 19, 21]) method to train networks of Krinskii-Kokoz neurons to mimic the low-dimensional attractors of different systems by using subthreshold voltage fluctuations and graded depolarization. This was accomplished by considering an approximation to the network structure that emerges when the voltages fluctuate near a resting membrane potential (*V*_*i*_ ≈ *V*_*R*_), the conversion of conductance-based synapses to current-based synapses:

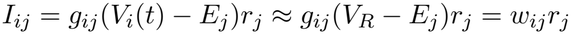

The weight matrix ***w*** was perturbed by a low-rank feedback term:

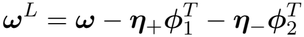

where ***ω***^*L*^ is the learned weight matrix after FORCE training. The low rank terms are functions of ***η***, a randomly generated *N* × *k* vector (commonly called the encoder), and ***ϕ*** a learned *N* × *k* vector called the decoder (see Materials and Methods for ***η***_*±*_ and ***ϕ***_1*/*2_ definitions. The matrix ***ϕ*** decodes an approximation to the *k*-dimensional supervisor ***x***(*t*) with

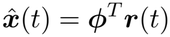

where ***r***(*t*) are the synaptic conductance gates.

The decoder was learned through the recursive least squares algorithm (Materials and Methods) which is an online optimization algorithm that can minimize the loss:

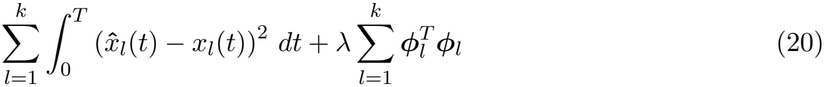

We successfully trained networks of Krinskii-Kokoz neurons with the current-based simplification to the synaptic filters to mimic various dynamical systems, all without inducing spiking (Figure 3). This included oscillators powered by stable subthreshold voltage fluctuations (Figure 3A-C), integrators (Figure 3D-E), which serve as a model for 1D path integration [46], pattern reproduction in the form of the notes of a song (Figure 3F), and low-dimensional chaotic attractors (Figure 3G). For all networks considered, not a single spike was fired during the training or testing regimes, with the only spikes generated in the initialization of the simulation.

**Figure 3.**
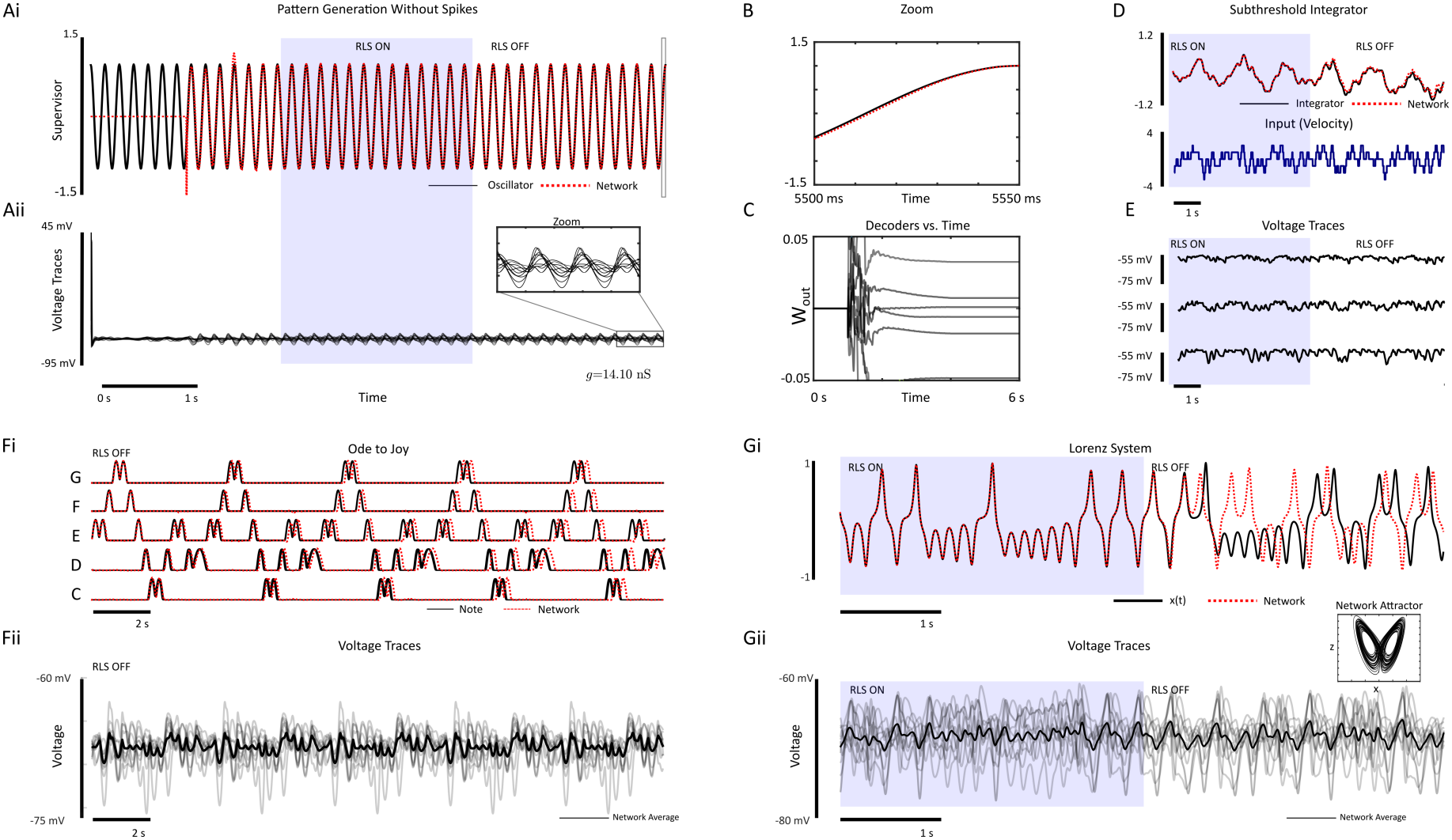
Subthreshold pattern generation in biophysically detailed networks without spikes. **(A)** A network of Krinskii-Kokoz neurons is trained with FORCE to mimic the dynamics of an oscillator with a period of 150 ms. (i) RLS is turned on after 2 seconds (blue panel), and turned off at the 4-second mark. The network’s approximant is shown in red while the target dynamics are in black (ii) The network’s subthreshold fluctuations are stabilized to produce the oscillator’s dynamics even with RLS turned off. B A zoom of the oscillator vs. approximant. **(C)** The decoders, ***ϕ*** vs time for 10 decoders. **(D)** A network trained to integrate an input current (blue, bottom) as a model for path integration. **(E)** The voltage fluctuations for 3 neurons (time aligned to (D)). **(F)** (i) A network trained to reproduce a pattern of activations corresponding to a sequence of notes for the song Ode to Joy. (ii) The voltage traces for 5 neurons. Note that no spikes are fired to reproduce the song. **(G)**(i) A network of Krinskii-Kokoz neurons was trained to reproduce the Lorenz attractor, a low-dimensional chaotic system. The network attractor is shown as an inset. (ii) The voltage traces for 5 neurons (time aligned to (G)). The network averages are shown in dark black.

The subthreshold fluctuations produced by voltage chaos were readily utilized for subthreshold computations by learning weights between neurons that produce stable subthreshold dynamics without spikes. However, we remark that spiking can be elicited with sufficiently strong feedback (Supplementary Figure 2) by drawing the encoder matrices ***η*** from wider distributions. This produces sparse spiking which does not destabilize the underlying computation (Supplementary Figure 2) while stronger feedback can destabilize computation (not shown).

### T-Type Calcium Channels are a Candidate for Graded Release Powered Sub-threshold Computations

The key assumption made in this modelling work was that the threshold for synaptic release could be dropped to near the resting membrane potential in the synaptic gating function:

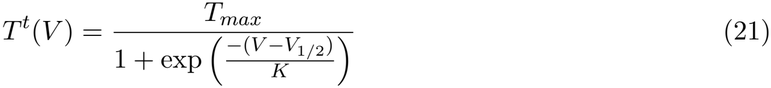

This corresponds partially to the low-voltage activated T-type calcium channel (Figure 4A). These channels operate in a regime near the resting membrane potential, but differ slightly from the release model considered here in Equation (21). T-type calcium channels activate and inactivate at low voltages which leads to a “window-current” that operates in a ≈ 20 mV interval near -60 mV. [13].

**Figure 4:**
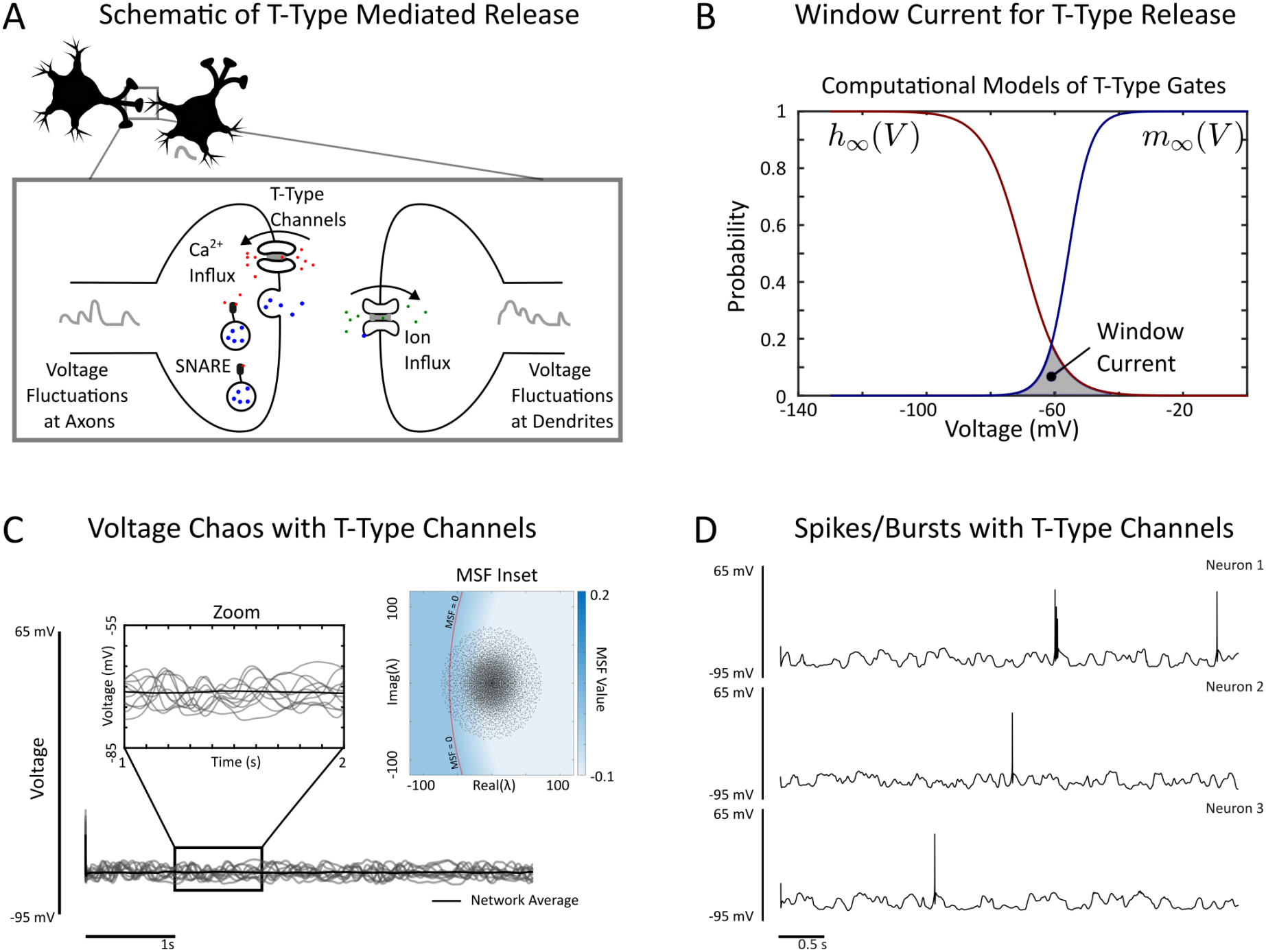
**(A)** A schematic of a putative candidate powering subthreshold voltage chaos: the T-type calcium channel. T-type calcium channels operate at low voltages near the resting membrane potential. They cause an influx of calcium, which can trigger vesicle release in a graded manner. Presynaptic voltage fluctuations would then lead to post-synaptic voltage fluctuations through T-type mediated graded depolarization. **(B)** A computational model fo the window current in T-type channels. The voltage activation (blue) and inactivation (red) lead to a window current operating in a narrow subthreshold voltage regime. **(C)** Voltage chaos emerges when incorporating a biophysical model of the T-type window current. The stability of the equilibrium state can be readily computed (inset). **(D)** For stronger conductances, spikes and bursts emerge in the voltage dynamics as in Figures 1 and 2.

We sought to determine if a more accurate model of the T-type channel’s window current could suppose subthreshold voltage chaos. First, we considered a modification of the synaptic gating function *T* (*V*) to more accurately model the window current:

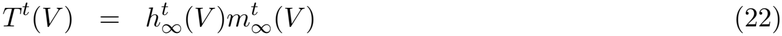

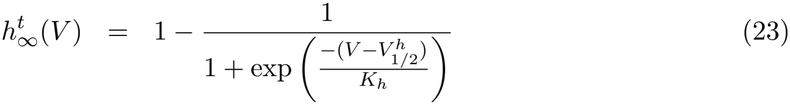

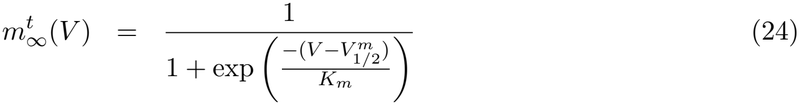

The functions 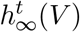 and 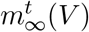 model the steady state inactivation dynamics and activation dynamics of the T-type calcium channel (Figure 4B). As in previous models of synaptic conductance dynamics [28], we assume that the influx of calcium, vesicle fusion, and neurotransmitter release is a suitably fast process that can be regarded as instantaneous for simplicity. This leads to postsynaptic conductance changes determined by the equation:

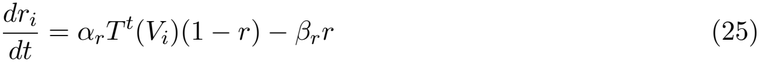

Next, we computed the Master Stability Function for networks of Hodgkin-Huxley neurons with the conductance gating dynamics given by equations (22)-(24) (Figure 4c). For strong enough conductances, the resting membrane potential destabilized and lead to robust and sustained voltage fluctuations (Figure 4c) without spikes. For stronger conductances, sparse spiking and bursting readily emerge (Figure 4D.

Finally, we investigated the impact of the window current width was on the ability of a network to generate subthreshold voltage fluctuations by varying the window parameters 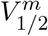 ,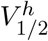 in addition to 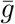 (Supplementary Figure 3). We subsequently measured the minimal spectral radius of the weights *w*_*ij*_ required to destabilize the stable resting membrane potential. We found that if 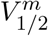 is too far away from ≈ *V*_*R*_ + 5 mV, the network would require stronger coupling (via 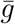) to destabilize the resting membrane potential. For the inactivation gate, we found that if 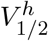 is sufficiently large, then *h*_*∞*_(*V*) ≈ 1 for any voltage *V* in the subthreshold range. However,if 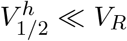 , then the window current would cease to exist, entirely, which also eliminates the subthreshold voltage fluctuations.

These results have shown that rich subthreshold dynamics emerge in biophysically detailed networks of neurons, and these subthreshold dynamics can be readily used for computation. Computational evidence supports the potential for low-voltage activated Calcium channels as mechanistically causing these dynamics in both *in vivo* and *in vitro* circuits.

## Discussion

Natural selection often drives circuits to find efficient solutions to specific problems to prevent metabolic energy from being needlessly wasted. The generation of spikes is metabolically costly, with each spike itself estimated to require 10^8^ − 10^9^ molecules of ATP [2]. As a result, spiking consumes approximately half of an excitatory neuron’s metabolic energy when the neuron is firing at low rates, while maintaining the resting membrane potential is far more efficient, consuming about 13% of the cell’s metabolic energy [2]. Further, for small organisms, like many invertebrates, local computations without spikes may be more efficient as information need not propagate far.

We have demonstrated that some circuits may be able to compute entirely without spikes by relying on subthreshold voltage fluctuations with low-voltage activated calcium channels (T-type). By coupling the neurons with conductances that are strong enough to destabilize the resting membrane potential, but not strong enough to elicit spikes, networks of neurons can produce rich subthreshold dynamics similar to a classical asynchronous state. In simpler networks comprised of 2D neuron models derived from the Hodgkin-Huxley neurons and fast synapses, the subthreshold dynamics can be shown to reduce to a kind of recurrent neural network where the activation function for the neurons in the subthreshold recurrent network is the voltage gate defined by low-voltage activated channels. This is in contrast to the conventional mapping between a neuron’s firing rate curve (f-I curve) and the activation in recurrent neural networks [34, 35] and machine learning more broadly. We remark that our work is complementary to these earlier studies, and that multiple types of balanced circuit dynamics may simultaneously co-exist. We also showed that the rich subthreshold dynamics could be readily tamed onto low-dimensional subthreshold attractors with FORCE training, although other techniques are also amenable. When combined with biologically plausible learning algorithms, our work shows that these subthreshold fluctuations can encode and compute useful information.

There is ample evidence to support the existence of graded depolarization, facilitating an analog or hybrid digital-analog code. Many invertebrate neural circuits do not fire spikes or rarely fire spikes and transmit information in an analog manner or graded manner [3, 8, 9, 47]. The retina in vertebrate circuits features analog encoding in retinal bipolar cells [48], but with digital all-or-nothing spikes also fired by bipolar cells [10, 49].

The primary mechanism we propose for subthreshold computation is through graded release powered by T-Type channels [50]. T-type channels form a complex allowing for action-potential independent release of neurotransmitter with Syntaxin-1A [17], while the calcium influx can trigger plasticity through an interaction between T-Type channels and Calmodulin, eventually leading to *α*CamKII activation [51], in addition to complexes with low-voltage potassium channels [52–54]. Both experiments and our computational modelling here support the role of T-Type channels for rich-subthreshold dynamics. Surprisingly, subthreshold fluctuations can also be direclty transmitted with spikes [55]. When paired with appropriately chosen spike-timing correlations, subthreshold voltage fluctuations can stabilize otherwise large voltage fluctuations into an intermediate range [56].

Our findings provide compelling evidence for the mechanism behind computations in neural circuits with very low spike rates. While spikes remain the primary vehicle for long-range communication, and certainly there are recurrent circuits that directly use spikes for computation, we have shown that specific circuits, primarily those in invertebrates, can operate in a ‘silent’ analog regime, leveraging graded subthreshold dynamics to execute complex computational tasks. Crucially, the expression of low-voltage-activated calcium channels provides a biophysical substrate for recurrent neural network dynamics in the complete or partial absence of action potentials.

## Acknowledgments

We thank Claudia Clopath, Amanda Foust, David Dupret, Vitor Lopes-dos-Santos, and Yonatan Aljadeff for their insightful comments. WN is funded by an NSERC Discovery Grant (RGPIN/04568-2020), a Canada Research Chair (CRC-2019-00416), and the Hotchkiss Brain Institute through the Cumming Medical Research Fund.

## Materials and Methods

We considered networks of conductance based neurons with conductance-based synaptic coupling:

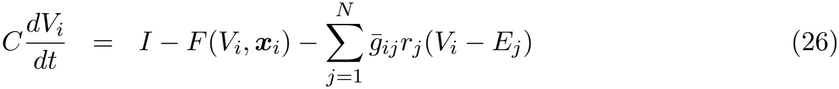

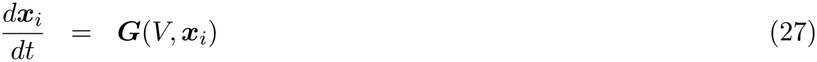

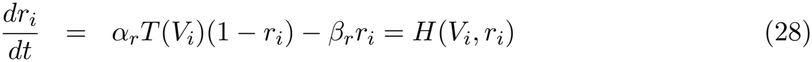

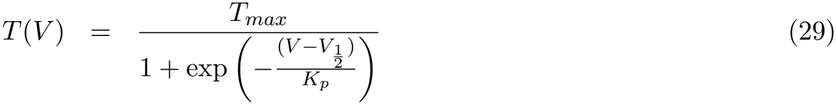

where 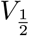 , the half-max synaptic release voltage was slightly more depolarized than the resting membrane potential *V* = *V*_*R*_, while all other parameters were the nominal parameters used for existing conductance based neurons. The voltage of a neuron is given by *V* (*t*) while the gating variables for various ionic conductances are collectively written as the vector ***x*** for generality. *C* acts as the neuronal capacitance, while the dynamics of *V* and the gating variables are given by the functions *F* (*V*, ***x***) and ***G***(*V*, ***x***), respectively.

We primarily considered the Hodgkin-Huxley neuron model:

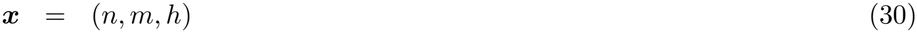

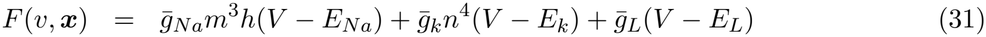

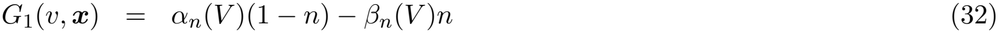

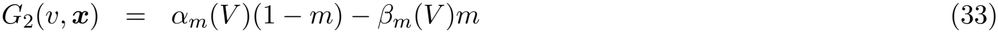

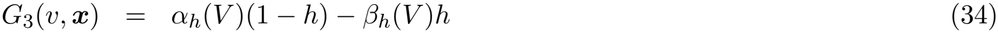

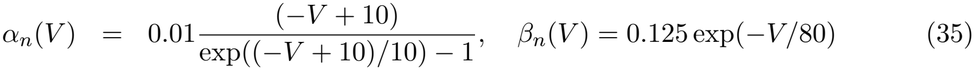

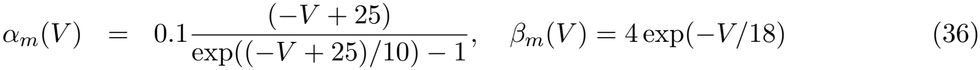

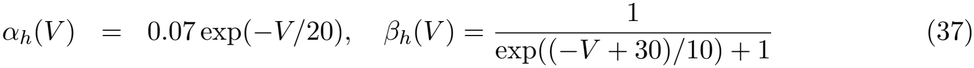

and the Krinskii-Kokoz neuron model:

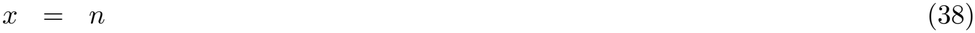

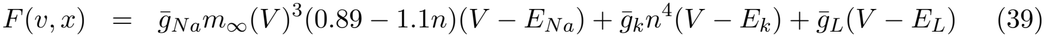

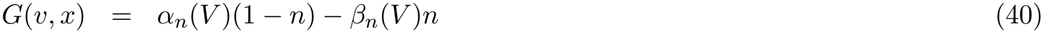

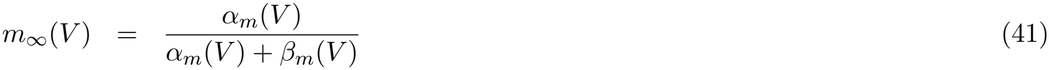

which is a 2-dimensional reduction of the Hodgkin-Huxley neuron models where all utilized gating functions are identical to the full Hodgkin-Huxley model. All neuron and network models were integrated with the Runge-Kutta45 integration scheme implemented in ODE45 in MAT-LAB2023A, with the exception of FORCE trained neurons which were integrated with a forward Euler method with Δ_*t*_ = 10^*−*2^ ms to avoid the complexities in adapting FORCE to work with RK45. We remark that the parameters listed in Table 1 and here correspond to the equilibrium potential zeroed to *V*_*R*_ = 0 as classically imposed by Hodgkin and Huxley [26]. The parameters (or equivalently y-axis of plots) can be shifted to a resting membrane potential of *V*_*R*_ = −65 mV. All connection parameters (*E*_*ampa*_/*E*_*gaba*_) and thresholds for synaptic release *V*_1*/*2_ are set assuming the same *V*_*R*_ = 0, but are rescaled for figures with *V*_*R*_ = −65*mV*.

### Loss of Stability of the Common Resting Membrane Potential

In order for a network of conductance neurons to have a common resting membrane potential *V*_*i*_ = *V*_*R*_, ***x***_*i*_ = ***x***_*R*_, the following conditions have to be imposed:

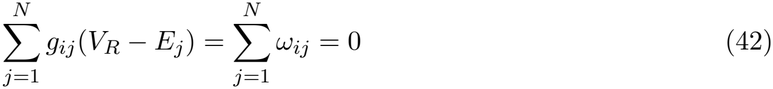

This allows for the derivation of the stability of the common resting membrane through the master stability function. In particular, if *λ*_1_, *λ*_2_, … *λ*_*N*_ are the eigenvalues of the weight matrix ***ω***, then the stability of the common resting membrane potential is identical to the stability of the **0** solution to the block system:

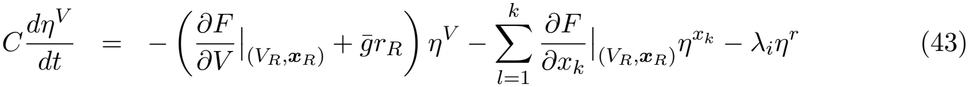

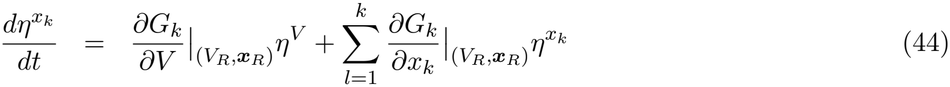

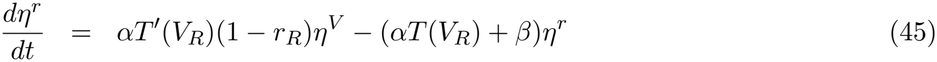

Where

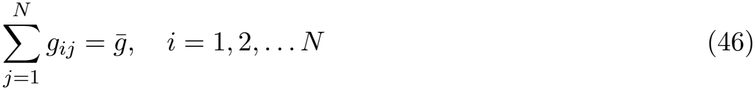

Each block consists of 2 + *l* equations with *N* total blocks where *l* is the number of gating variables. The eigenvalues of the Jacobian of the **0** solution in equations (91)-(93) above are functions of the eigenvalues (*λ*_*i*_) of ***ω***. Defining the eigenvalues of the Jacobian as *μ*, then the master stability function is defined as the maximum real component of an eigenvalue *µ*for all *N* blocks. As the blocks are functions of *λ*, the master stability function can be computed though discretizing *λ* over a 2D mesh in the complex plane, with the eigenvalues *µ*of equations (91)-(93) numerically computed over the 2D mesh. Then, if any eigenvalue *λ*_*i*_ lies in the region where the master stability function is positive, the common resting membrane potential *V*_*i*_ = *V*_*R*_, ***x***_*i*_ = ***x***_*R*_ has lost stability. Further, in the supplementary information, we show a case where the onset of instability can be computed exactly for 2D neuron models with fast synapses.

### First Order Reduced and Controlled Error (FORCE)

The first order reduced and controlled error (FORCE) technique was applied to learn a matrix of *N* × *k* decoders ***ϕ***, that decode out the dynamics of some supervisor *z*(*t*) with 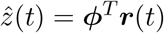 where ***r***(*t*) is the vector of synaptic conductance gates given by *r*_*i*_(*t*) = *r*_*∞*_(*V*_*i*_(*t*)). Due to their numerical efficiency and ease of applicability, we considered networks of Krinskii-Kokoz neuron’s with current based synapses:

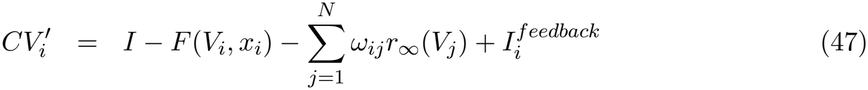

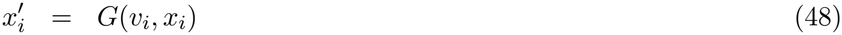

were 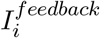 is the learned feedback generated by FORCE training. The current-based synaptic approximation is appropriate when the neurons voltage is near the resting membrane potential. The decoders are learned with recursive least squares (RLS):

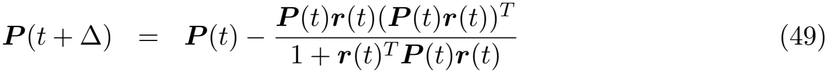

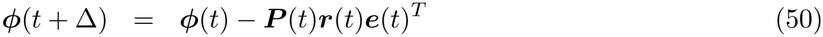

where 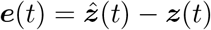.

RLS minimizes the squared loss: 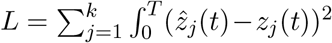 over the interval [0, *T*] (when RLS is on). A regularization term to the loss is added by initializing ***P*** (0) = ***I***_*N*_ *α*^*−*1^, as ***P*** acts as a rolling estimate of the inverse of the Grammian matrix 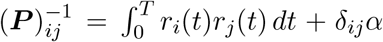 where *d*_*ij*_ is the Kroenecker delta function.

To stabilize the dynamics of the network onto the supervisor, while preserving Dale’s law, the decoder is not directly fed back into the network like in conventional applications of FORCE training. Instead, the “encoders”, which are randomly generated from a uniform distribution on [-Q,Q] are decomposed into positive and negative components as in [57]:

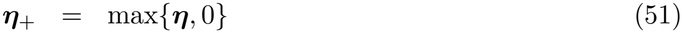

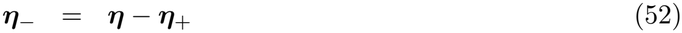

The decoder learned from RLS can be written as ***ϕ*** = [***ϕ***^*E*^, ***ϕ***^*I*^] where *E* and *I* correspond to the excitatory and inhibitory subpopulations. Then, the feedback to the network is given by:

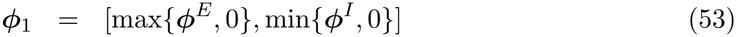

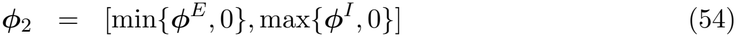

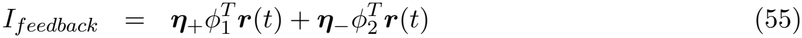

and the resulting low-rank perturbations lead to the resulting overall weight matrix

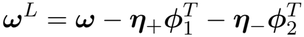

which satisfies Dale’s Law, where ***ω***^*L*^ is the learned weight matrix [57]

### Additional Methods for Each Figure

#### Figure 1

The network consisted of *N* = 8000 excitatory neurons and *N* = 2000 inhibitory neurons. The excitatory/inhibitory nature of the neurons was imposed through the reversal potential of the synapses they formed, either *E*_*gaba*_ for inhibitory neurons or *E*_*ampa*_ for excitatory neurons. In order to compute the master stability function, the resting membrane state was computed by solving *F* (*V*, ***x***) = 0 and ***G***(*V*, ***x***) = 0 for *V*_*R*_, ***x***_*R*_ by using MATLAB’s solver *fsolve*. The equilibrium point is necessary for computing the coefficients of the master stability function blocks (Supplementary Derivation 1). The network was sparsely coupled with a sparsity of *p* = 0.005. In order to generate a random matrix which satisfies the conditions 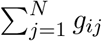 and 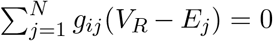, we first wrote the constraints as a linear system for 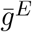 and 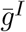, which is the row sum of all the excitatory and inhibitory conductances. This yields:

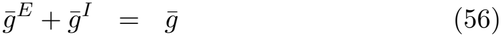

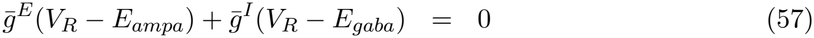

which is easily solvable for 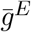 and 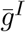:

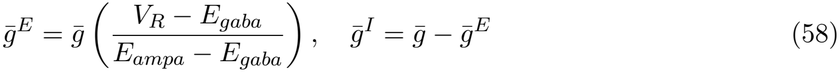

Then, a random matrix of conductances are drawn from the standard uniform distribution [0, 1] with a sparsity of 0.005. The first *N*^*E*^ neurons are treated as excitatory with the last *N*− *N* ^*E*^ neurons are treated as inhibitory. The mean of the first *N* ^*E*^ conductances is set to 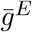 for each neuron by dividing each element by the empirical mean, and multiplying by 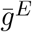, with a similar operation used for the last *N* − *N* ^*E*^ elements with 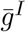. This procedure ensures the generation of a matrix of conductances that satisfies both the constant conductance and weight balance conditions.

#### Figure 2

We considered networks of 2000 excitatory and 2000 inhibitory neurons, with a sparsity of 0.01. The matrix of conductances were generated as in Figure 1. The Gaussian distribution and exponential distribution were fit by empirically measuring the mean and standard deviation of the voltage fluctuations (Gaussian), and the mean inter-spike-interval (exponential), as both distributions are defined by only these parameters, respectively.

#### Figure 3

For the oscillator (Figure 3(A-C), 250 excitatory and 250 inhibitory neurons were used, with the elements of the encoders ***η*** being generated from a uniform distribution on [−3, 3]. The alpha parameter was set to 10 and RLS was applied every 20 time steps. The supervisor was

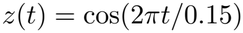

where the period is 0.15 seconds. As current based synapses were used, the weight matrix *ω* was directly generated with a sparsity level of *p* = 0.05, with the first *N*^*E*^ components of each row being either 0 (with probability 0.95) or 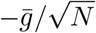 (with probability 0.05), and the last *N*^*I*^ components being either 0 or 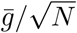 , with 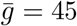.

For the integrator (Figure 3D-E), A network of 500 excitatory and 500 inhibitory neurons with current based synapses was trained to learn to integrate inputs. A random step-signal was generated with:

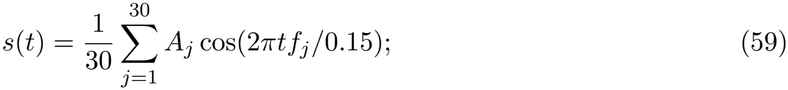

where *f*_*j*_ was randomly generated from a standard uniform distribution and *A*_*j*_ was randomly generated from a standard normal distribution. The input *c*(*t*) was then generated by Z-transforming *s*(*t*) and subsequently rounding the result to the nearest integer at each point in time. The supervisor for the network, *z*(*t*) was computed as as

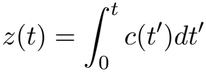

A total of 19 seconds of training time (RLS on) was used after the initial second of simulation with RLS off. The *α* parameter was taken to be 10, while RLS was applied every 2 time steps. Weights were identically generated as in the integrator example.

The Lorenz network was identical to the oscillator network, only the encoder elements generated from the uniform distribution from [−2, 2] and with 1500 excitatory and 1500 inhibitory neurons. RLS was applied every 2 time units, with *α* = 2. The Lorenz system is given by

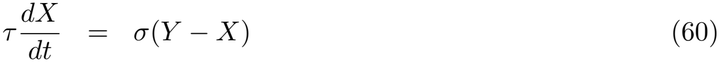

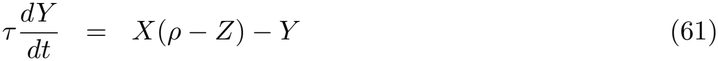

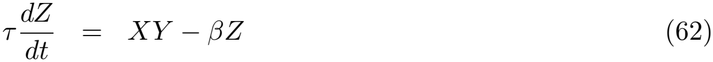

with *s* = 10, *ρ* = 28, and *β* = 8*/*3 where *τ* = 200 parameterizes time to operate on the time-scale of neurons, while each component *X*(*t*), *Y* (*t*), *Z*(*t*) is z-transformed before being applied as a supervisor. RLS was applied every 2 time units. A total of 30 seconds of training was used.

For the sequences of notes corresponding to the first bar of Ode to Joy, we utilized a high-dimensional temporal supervisor as in [21]. Additional details on how the supervisor was constructed can be found in [21] along with a copy of the supervisor in the accompanying code. All supervisor components were smoothed with a 50 millisecond smoothing window to prevent oscillations in the conductance network near the onset/offset of a note. A total of 2000 excitatory and 2000 inhibitory neurons were used, with identical parameters as in the oscillatory supervisor (Figure 3A).

#### Figure 4

We considered networks of 4000 excitatory neurons and 1000 inhibitory neurons. The conductances were generated as in Figure 1, with a sparsity of 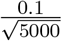 . The parameters for the gating function were 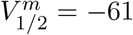 mV (4 mV with 0 mV resting), 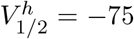 (-10 mV with 0 mV resting), and *K*_*m*_ = 3.5 mV, *K*_*h*_ = 6 mV. The conductances for the Figure 4C was 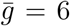 while for 4D was 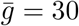 nS. The bias current *I* was lowered to *I* = −5 pA for these simulations.

## Supplementary Information

### Supplementary Derivation 1: Master Stability Function for Conductance Based Synapses

Conductance-based neuron models, originally proposed by Hodgkin and Huxley are biophysically detailed neuron models consisting of a voltage term *V* (*t*) and a number of gating variables that we will represent generally with the vector ***x***(*t*). The dynamics of *V* (*t*) and ***x***(*t*) are given by the equations:

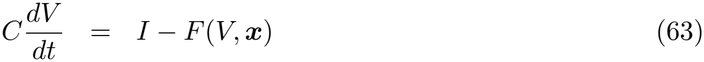

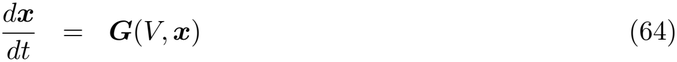

where *C* is the membrane capacitance, and *I* is an applied current. The vector of gating variables ***x***(*t*) collectively control the spike-generation, integration, and subthreshold dynamics of the neuron, along with the dynamics of *V* (*t*) through *F* (*V*, ***x***). The gating dynamics are determined by the multi-variable function ***G***(*V*, ***x***).

We will make the assumption that the single neuron model in (63)-(64) operates in the stable-resting membrane potential regime, where *I < I*^*\**^ is less than some bifurcation point of the system (63)-(64) where stable spiking emerges for *I > I*^*\**^. In particular, the neurons are constrained to have a stable resting membrane potential (*V* = *V*_*R*_) with associated steady state gating variables ***x*** = ***x***_*R*_.

The neurons are coupled through conductance-based synaptic coupling given by:

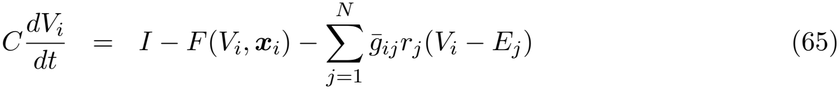

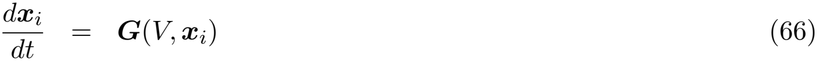

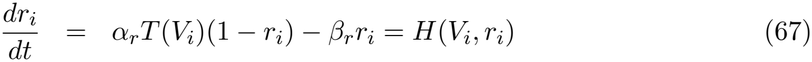

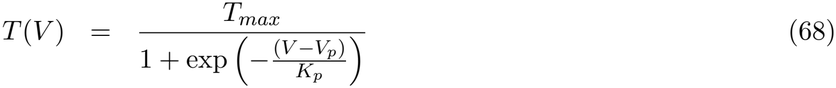

Equations (65)-(68) form a coupled network of conductance-based neurons with conductance-based synapses. Next, we will assume that the voltage for each neuron fluctuates near (*V*_*R*_, ***x***_*R*_), which also leads to an equilibrium solution for *r, r*_*R*_:

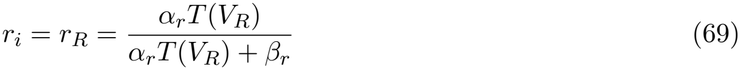

At the network level,the solution *V*_*i*_ = *V*_*R*_, ***x***_*i*_ = *x*_*R*_, ***r***_*i*_ = *r*_*R*_ exists for *i* = 1, 2, … *N* if

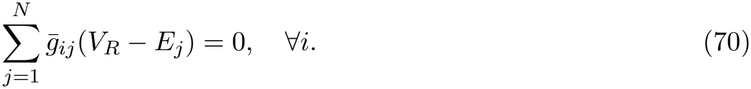

Equation 70 can be interpreted as a “precise balance” condition if we consider excitatory (*V*_*R*_ *< E*_*j*_) and inhibitory (*V*_*R*_ *> E*_*j*_) synapses, which is immediately satisfied so long as

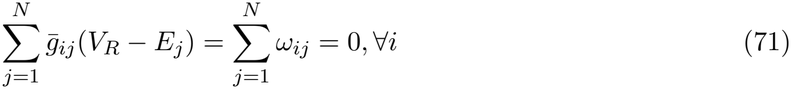

Next, the network’s fluctuations are assumed to occur near the resting membrane potential for all neurons by considering the perturbations (***ϵ***^*V*^ , ***ϵ***^***x***^, ***ϵ***^*r*^) with: 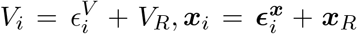,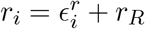 for *i* = 1, 2 … *N* . Expanding out and linearizing yields:

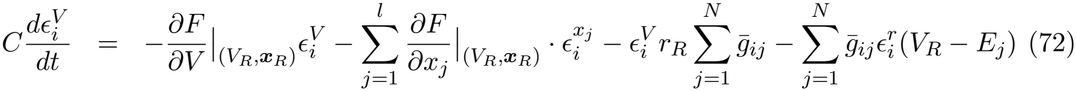

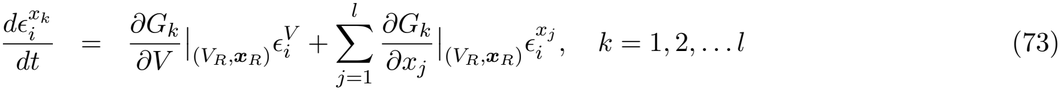

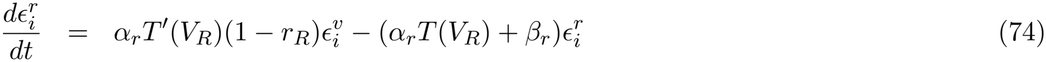

Next, we will make an additional simplifying assumption: 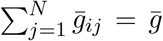 is a constant, which leads to the following system:

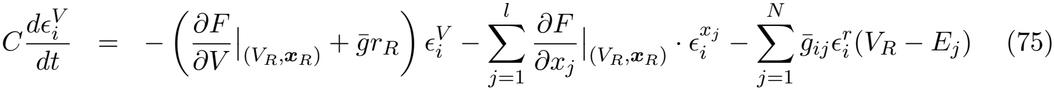

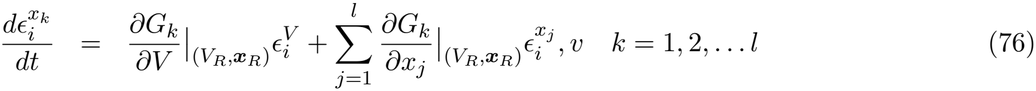

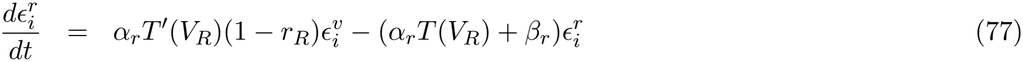

Which can be written in block matrix form as the following:

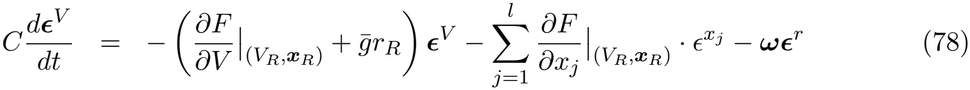

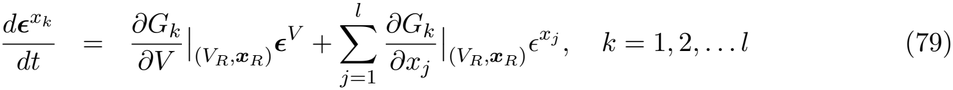

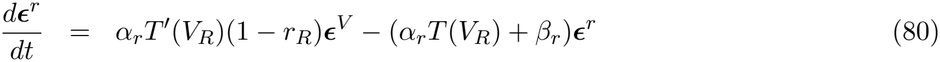

Before proceeding further, we remark that the two following conditions were imposed on the conductances:

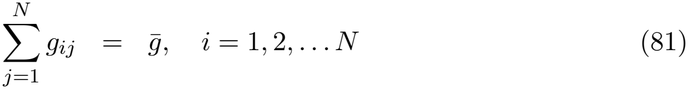

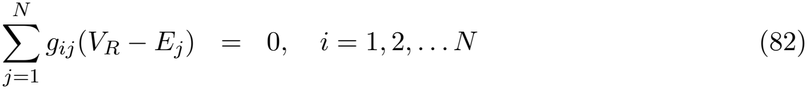

Neither condition is necessary for the existence of rich subthreshold dynamics. However, both conditions are extremely convenient mathematically to determine the stability (or lack thereof) of the stable-resting membrane potential *V* = *V*_*R*_, ***x*** = ***x***_*R*_, and the potential emergence of rich subthreshold states.

In total, with *l* gating variables, a conductance network of *N* neurons leads to (2 + *l*) × *N* variational equations. The eigenvalues of the variational system (91)-(93) determine the stability of the global resting membrane solution. Despite the the complex nature of these equations, they simplify greatly as the coefficients in front of the vectors ***ϵ***^*V*^ , 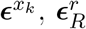, are either scalars, or the matrix ***ω***. We can exploit the latter fact by assuming further that the matrix ***ω*** is diagonalizable

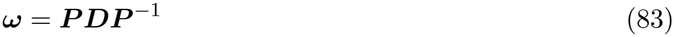

where ***D*** corresponds to the diagonal matrix of eigenvalues of *ω* and ***P*** is the matrix eigenvectors of ***ω*** and considering the substitutions

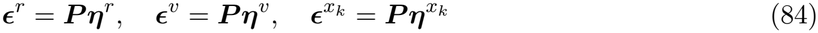

which yields:

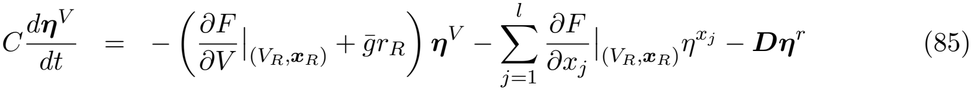

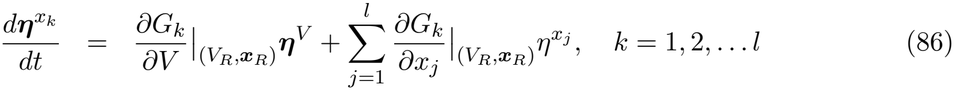

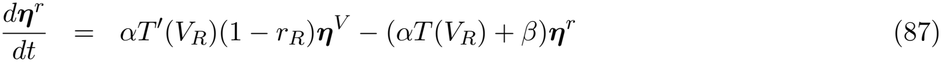

which is a block-diagonal linear system corresponding to:

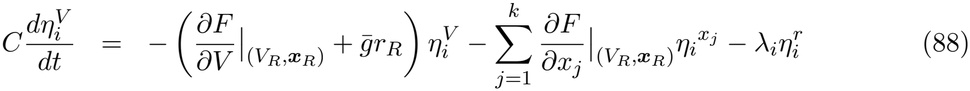

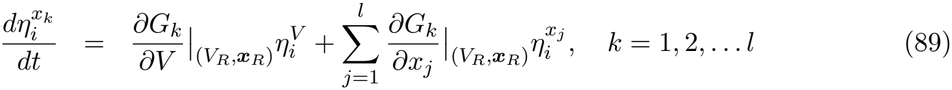

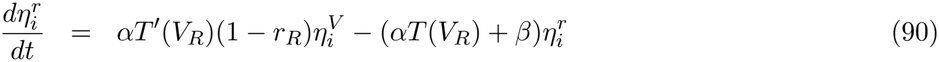

where *λ*_*i*_ is an eigenvalue of ***ω***.

Through the Master Stability Function framework, as the blocks are identical, they can all be replaced with a generic block that is a function of a *λ* parameter which acts as a stand in for any eigenvalue of ***ω***:

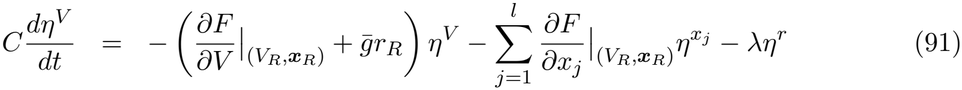

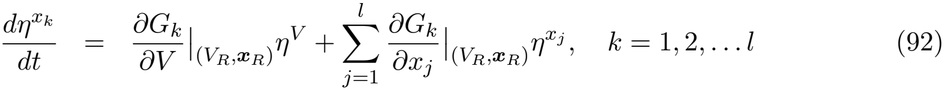

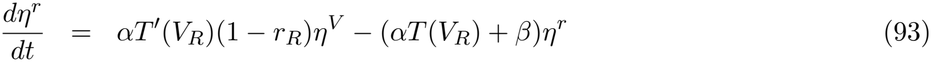

The resulting system, leads to *l* + 2 eigenvalues for the equilibrium 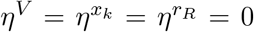 as a function of *λ, μ*_1_(*λ*), *μ*_2_(*λ*), … *μ*_2+*l*_(*λ*) with the master stability function (MSF) defined as

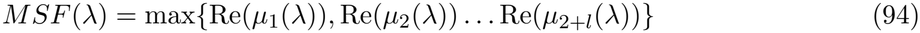

The equilibrium *V*_*i*_ = *V*_*R*_, ***x***_*i*_ = ***x***_*R*_, *r*_*i*_ = *r*_*R*_ loses stability when the real component of the master stability function is positive for some eigenvalue of ***ω***:

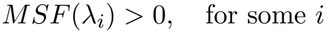

### Supplementary Derivation 2: An Exactly Solvable Case: 2D Neuron Models with Fast Synapses

For the case where 1) the neuron model in question has two dimensions, and 2) the synapses are fast, we can consider an approximation that yields a functional form for the onset of instability of the resting membrane potential. Consider the following:

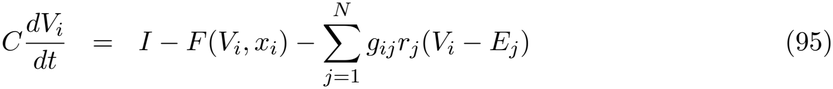

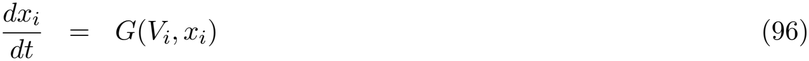

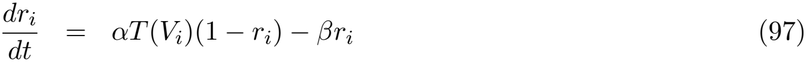

IF the synapses are faster than the voltage dynamics, the synaptic gating variables can be taken to be:

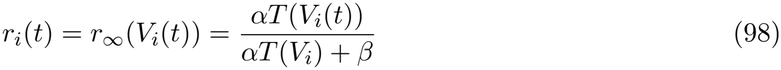

which yields the following reduced 2*N* dimensional system:

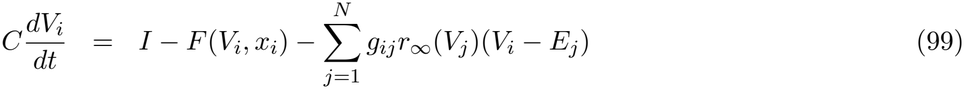

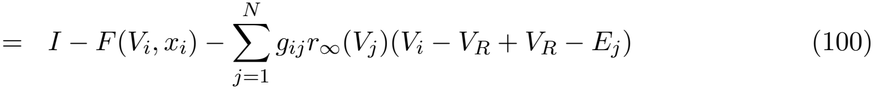

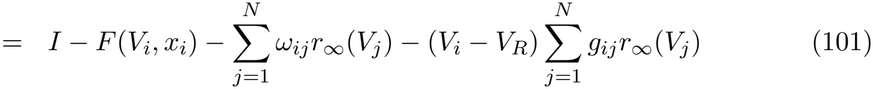

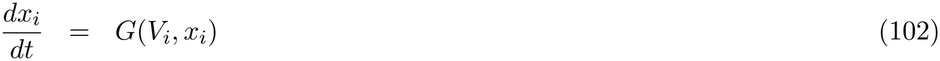

and we will assume the same conditions as in the generic conductance-based case considered above. Expanding around the common resting membrane state with 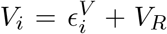 and 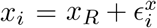 yields:

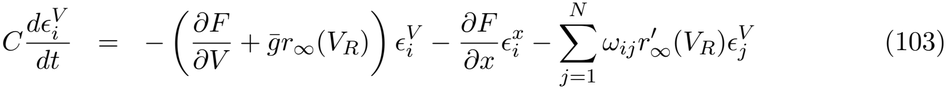

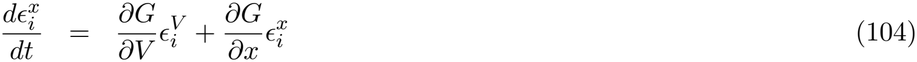

which yields the following set of equations used to determine the master stability function:

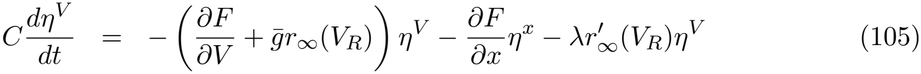

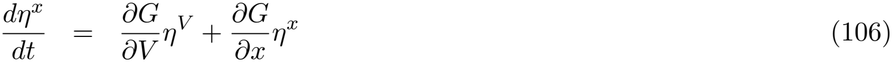

Suppose *μ*(*λ*) is an eigenvalue of the Jacobian of *η*^*V*^ = 0, *η*^*x*^ = 0 as a function of *λ*, then with the following three conditions:

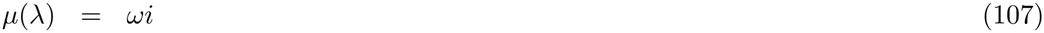

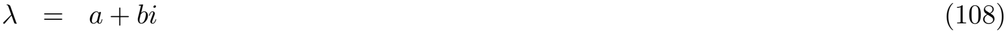

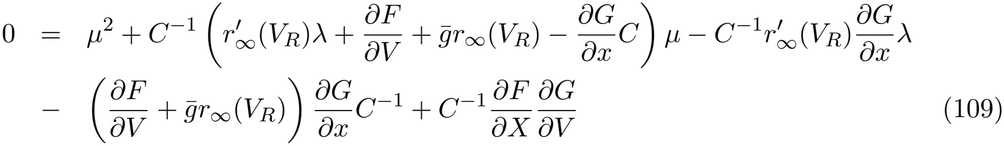

yields the following equation describing the curves that yield a master stability function of 0 (loss of stability of *V*_*i*_ = *V*_*R*_, *x*_*i*_ = *X*_*R*_):

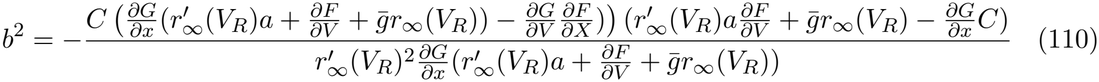

### Supplementary Derivation 3: Reduction of 2D Neuron’s with Fast Synapses to Adapting Rate Networks

Upon expansion of 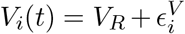 and 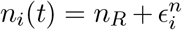 for the 2D Krinskii-Kokoz network with fast synapses, the system becomes:

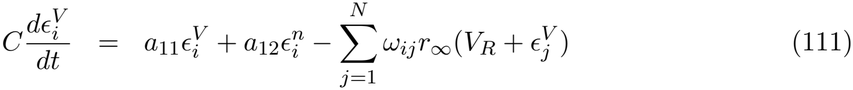

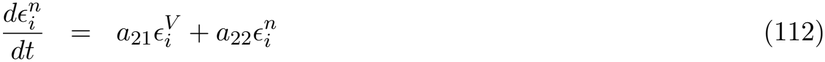

where we have partially linearized the subthreshold dynamics, but have left 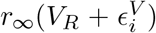 as nonlinear. The coefficients are given by

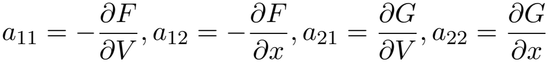

The partial linearization is justified by the fact that the resting state exists for the isolated neuron *V*_*R*_, *n*_*R*_ and thus if the fluctuations are near the resting state, the only term that is significantly non-linear is *r*_*∞*_(*V*). Next, a rescaling of time and space with:

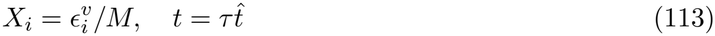

yields:

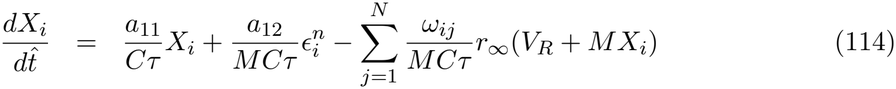

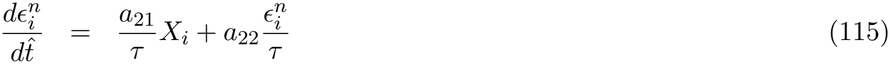

which leads to the following set of transforms/parameterizations

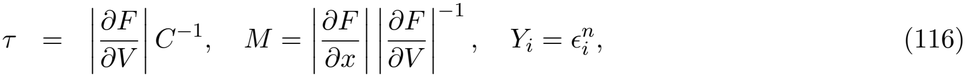

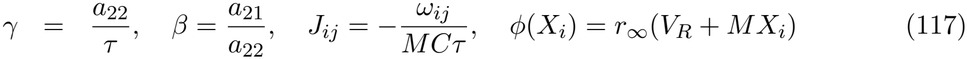

that converts equations (114)-(115) to

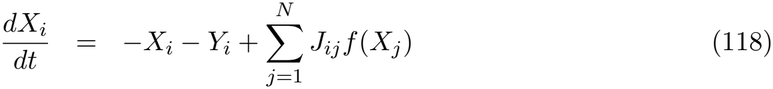

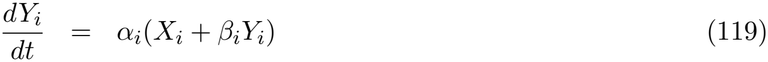

which is the rate network considered in [33].

## Supplementary Figures

**Supplementary Figure 1:**
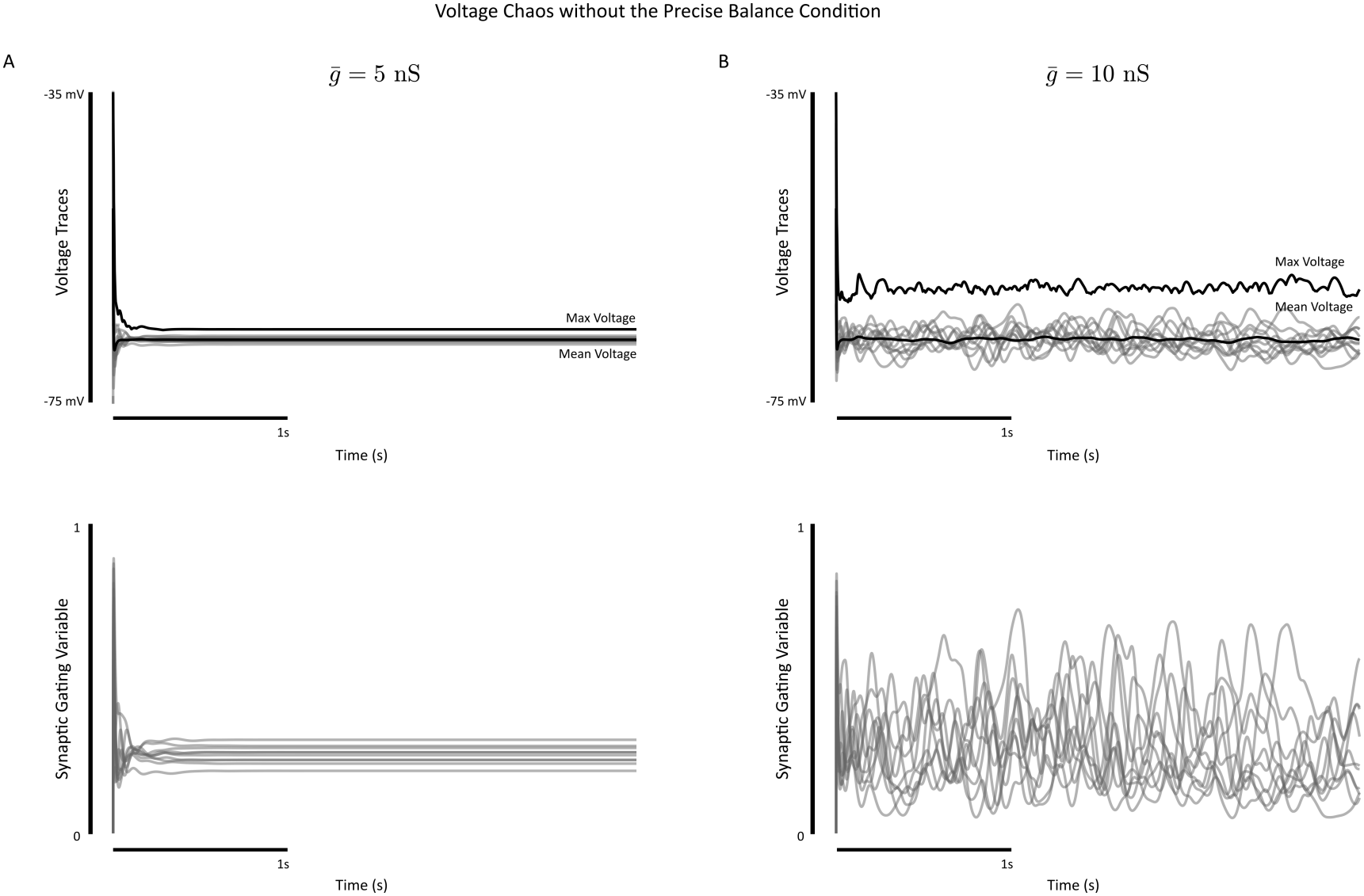
Voltage chaos without precise balance. A network of 8000 excitatory neurons and 2000 inhibitory Hodgkin-Huxley neurons is simulated for 3 seconds with 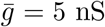 (left) and 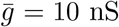 (right). **(A)** The voltages for 10 neurons (top) and conductance gating variables *s*(*t*) (bottom). The max and mean voltages are plotted in dark black for all neurons in the network, indicating no spiking has taken place. **(B)** Identical as in (A). The conductances were generated as in Figure 1, with precise balance and then perturbed with 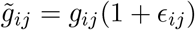 where *ϵ*_*ij*_ is a uniform random variable on [0, 0.2].

**Supplementary Figure 2:**
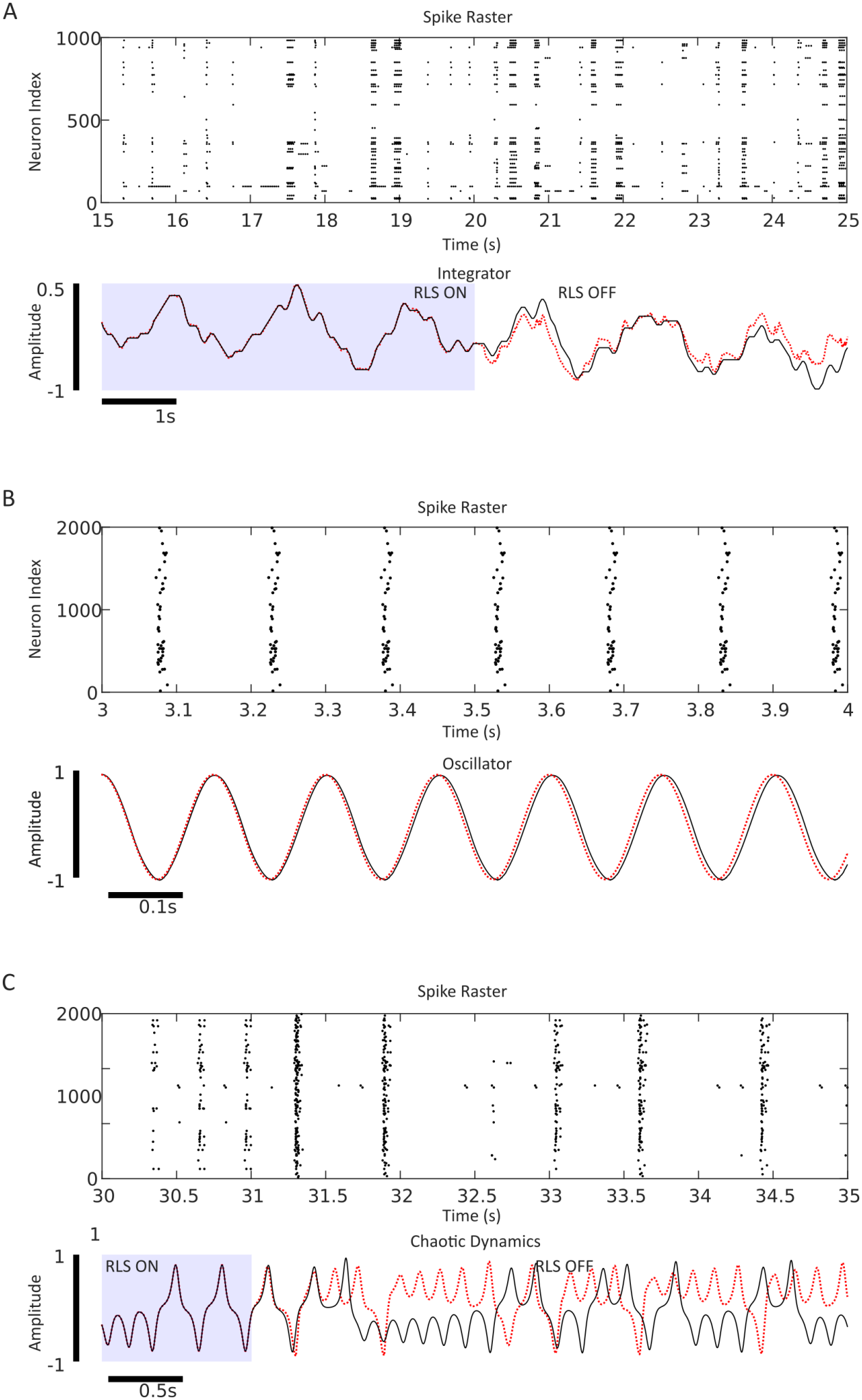
Sparse spiking and subthreshold dynamics enable computations. **(A)** A network identical to the one considered in Figure 3D was trained to integrate a velocity like input. Sparse spiking was elicited by generating the encoder elements ***η***_*i*_ from a wider distribution, *η*_*i*_ ∈ [−5, 5]. (Top) The spike raster plot for all neurons is shown. (Bottom) The network (red dashed line) vs. supervisor (black solid line). The network average firing rate during testing was 0.00017 Hz. **(B)** A network identical to the one considered in Figure 3A was trained to oscillate. Sparse spiking was elicited by generating the encoder elements ***η***_*i*_ from a wider distribution, *η*_*i*_ ∈ [−10, 10]. (Top) The spike raster plot for all neurons is shown. (Bottom) The network (red dashed line) vs. supervisor (black solid line). The network average firing rate during testing was 0.000015 Hz. **(C)** A network identical to the one considered in Figure 3G was trained to display a chaotic attractor. Sparse spiking was elicited by generating the encoder elements ***η***_*i*_ from a wider distribution, *η*_*i*_ ∈ [−8, 8]. (Top) The spike raster plot for all neurons is shown. (Bottom) The network (red dashed line) vs. supervisor (black solid line). The network average firing rate during testing was 0.00004 Hz.

**Supplementary Figure 3:**
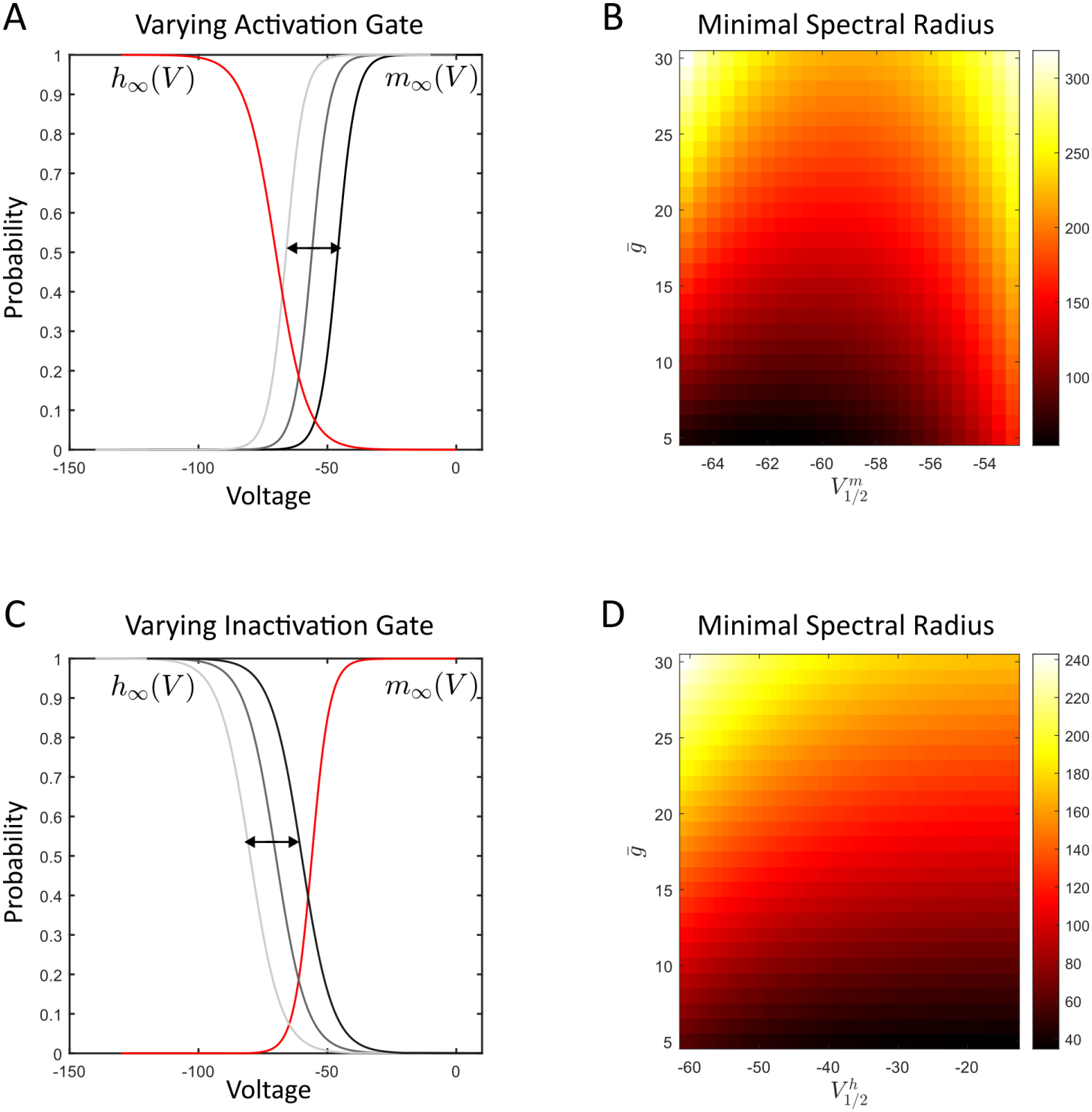
The impact of the inactivation and activation gates in T-type calcium channels on subthreshold voltage chaos. **(A)** The steady state activation gate *m*_*∞*_(*V*) is shifted through the 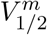 parameter. **(B)** The minimal spectral radius of the weight matrix required to destabilize the common resting membrane potential (brighter colours denote larger values) as a function of 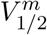 and 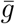. Note the non-monotonic effect where the smallest spectral radius required to destabilize the resting state occurs when 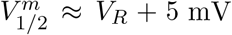 mV. **(C)** The steady state inactivation gate *h*_*∞*_(*V*) is shifted through the 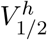 parameter. **(D)** The minimal spectral radius of the weight matrix required to destabilize the common resting membrane potential (brighter colours denote larger values) as a function of 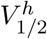 and 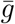. For larger values of 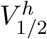 , the inactivation gate has no effect on the spectral radius as *h*_*∞*_(*V*) ≈ 1 for all subthreshold voltages.

